# A vagal sensory–hypothalamic oxytocin–brown adipose tissue pathway mediates cholecystokinin-induced thermogenesis

**DOI:** 10.1101/2025.08.11.669026

**Authors:** Yuta Masuda, Miyuki Kawase, Rika Kitano, Kento Ohbayashi, Toshiki Yabe-Wada, Hiroshi Inoue, Mamoru Tanida, Yusaku Iwasaki

**Author notes:** These two authors contributed equally to this work (Yuta Masuda, Miyuki Kawase). Corresponding authors **Dr. Mamoru Tanida** Department of Physiology II, Kanazawa Medical University, 1-1, Daigaku, Uchinada, Ishikawa 920-0292, Japan, **Prof. Yusaku Iwasaki** Laboratory of Animal Science, Graduate School of Life and Environmental Sciences, Kyoto Prefectural University, 1-5 Hangi-cho, Shimogamo, Sakyo-ku, Kyoto 606-8522, Japan.

## Abstract

Thermoregulation is essential for survival in homeothermic animals. Vagal sensory nerves are well known to detect visceral signals and regulate physiological functions, including feeding, metabolism and immunity. However, their role in thermoregulation remains poorly understood. Cholecystokinin (CCK), a gut hormone released postprandially, activates vagal sensory nerves via CCK-A receptors (CCK-AR). Exogenous CCK has been reported to induce thermogenesis in the intrascapular brown adipose tissue (iBAT), but the involvement of vagal sensory nerves and the central neural mechanisms that mediate this effect are not fully understood. In this study, we assessed the thermogenic effect of intraperitoneally (I.P.) administered CCK-8 and investigated the underlying autonomic reflex pathways. I.P. CCK-8 transiently and dose-dependently increased rectal temperature. This response was significantly attenuated by pharmacological blockade of CCK-AR, subdiaphragmatic vagotomy, or knockdown of CCK-AR primarily targeting in vagal sensory neurons. In addition, CCK-8 activated sympathetic nerve activity via vagal afferents. CCK-8–induced thermogenesis was blunted by iBAT sympathectomy or β3-adrenergic receptor blockade. Furthermore, I.P. CCK-8 activated oxytocin neurons in the paraventricular nucleus of the hypothalamus (PVH^OXT^). Chemogenetic inhibition of PVH^OXT^ neurons or intracerebroventricular administration of an oxytocin receptor (OXTR) antagonist attenuated the thermogenic response. These findings demonstrate, for the first time, the full neural circuitry underlying CCK-induced thermogenesis by delineating its afferent input (CCK-AR-expressing vagal sensory neurons), central integrative hub (PVH^OXT^ neurons and OXTR signaling), and efferent output (iBAT sympathetic nerves). This study further suggests that CCK-AR -expressing vagal sensory neurons may contribute to thermoregulation under physiological conditions in which CCK is endogenously released.

**Key points summary:**

- Vagal sensory nerves, which connect the gut and the brain, play a key role in regulating meal-related physiology, however their role in thermoregulation remains incompletely understood.
- This study reveals for the first time the full autonomic reflex pathways underlying thermogenic effect of the gut hormone cholecystokinin (CCK), comprising afferent input (CCK-A receptor-expressing vagal afferents), a central integrative hub (oxytocin neurons in the hypothalamic paraventricular nucleus; PVH^OXT^ neurons), and efferent output (intrascapular brown adipose tissue via sympathetic nerves).
- Both CCK-A receptors-expressing vagal afferents and sympathetic nerves innervating brown adipose tissue are required for thermogenesis induced by exogenous CCK-8.
- Activation of PVH^OXT^ neurons by CCK-8 is critically involved in mediating this thermogenic effect.
- This newly identified gut-brain-fat axis may contribute to part of diet-induced thermogenesis, and its impairment could be involved in the development of metabolic disorders such as obesity.

## Introduction

Vagal sensory nerves connect the thoracoabdominal organs with the brain, sensing locally released factors via their peripheral terminals in a paracrine or synapse-like fashion and relaying this information to the central nervous system (Berthoud *et al*., 2025). These nerves regulate a wide range of physiological functions, including gastrointestinal and cardiovascular activity as well as immune responses (Paintal, 1973; Young *et al*., 1974; Pavlov & Tracey, 2012; Borgmann *et al*., 2021), and they play a particularly important role in controlling meal-related physiology such as feeding behavior and metabolic regulation (Mendez-Hernandez *et al*., 2025). Although the roles of vagal sensory nerves in feeding and glucose metabolism have been extensively studied (Krieger *et al*., 2016; Tan *et al*., 2020; Li *et al*., 2022; McDougle *et al*., 2024), their contribution to thermoregulation including postprandial thermogenesis remains poorly understood.

The activity of vagal sensory neurons is modulated by gastrointestinal and pancreatic hormones, which exhibit meal-related fluctuations in secretion (Iwasaki & Yada, 2012; Yada *et al*., 2025). Among these, cholecystokinin (CCK), a potent activator of vagal sensory neurons, is a key intestinal hormone released postprandially (Lankisch *et al*., 2002; Date *et al*., 2002; Simasko *et al*., 2002). CCK is secreted from enteroendocrine I-cells in the duodenum and jejunum, primarily in response to dietary fat and protein intake (Liddle, 1995). CCK exists in multiple peptide forms of different lengths, with the shortest bioactive form, CCK-8, being sulfated at the seventh tyrosine residue from the C-terminus (Dufresne *et al*., 2006). CCK acts on CCK-A receptors (CCK-ARs) expressed on vagal sensory neurons, transmitting signals to the nucleus tractus solitarius in the brainstem (Moran *et al*., 1992; Mönnikes *et al*., 1997). Previous studies have shown that peripheral administration of CCK-8 increases thermogenesis in intrascapular brown adipose tissue (iBAT), and this effect is abolished by either cervical or subdiaphragmatic vagotomy (Yamazaki *et al*., 2019; Wang *et al*., 2019). However, whether this thermogenic action of CCK is mediated via CCK-ARs expressed on vagal sensory neurons remains unclear. In addition, activation of vagal afferent pathway by CCK administration has been reported to stimulate hypothalamic oxytocin neurons in the paraventricular nucleus (PVH^OXT^ neurons) (Olson *et al*., 1992; Blevins *et al*., 2003), contributing to the mediation short-term feeding suppression (Xi *et al*., 2017). Furthermore, PVN^OXT^ neurons have also been shown to regulate thermogenesis in iBAT (Fukushima *et al*., 2022). Although the CCK–vagal afferents–PVH^OXT^ neuron axis has been implicated in feeding regulation, its potential involvement in thermogenesis has yet to be elucidated.

In this study, we aimed to determine whether the CCK–vagal afferents–PVN^OXT^ neuron axis regulates thermogenesis in iBAT and to elucidate its underlying neural mechanisms. We first examined whether intraperitoneal (I.P.) administration of CCK-8 increases rectal temperature in mice as an index of core body temperature. We then evaluated the involvement of CCK-AR on vagal sensory nerves using subdiaphragmatic vagotomy, pharmacological blockade, and knockdown of CCK-AR predominantly in vagal afferents. To investigate the efferent pathway, we assessed the roles of sympathetic nerves innervating iBAT and β3-adrenergic receptor signaling. Finally, we evaluated the contribution of PVH^OXT^ neurons and central OXT receptor signaling using a chemogenetic approach and pharmacological inhibitors. Our results demonstrate that CCK-8 increases core body temperature via activation of vagal afferents expressing CCK-AR, PVH^OXT^ neurons, central OTR signaling, and iBAT sympathetic nerves, thereby inducing thermogenesis in iBAT. These results provide new insight into the autonomic circuitry linking postprandial gut signals to thermoregulatory effectors.

## Materials and methods

### Ethical approval

All procedures were performed in accordance with the ethical guidelines of The Journal of Physiology and the Japanese Act on Welfare and Management of Animals. The experimental protocols were approved by the Institutional Animal Care and Use Committee of Kyoto Prefectural University (Approval Nos. KPU060327-RC-4, KPU060327-RC-5, KPU060327-C4).

### Animals

Wild-type C57BL/6J mice (The Jackson Laboratory Japan, Inc., Yokohama, Japan), Phox2b-Cre mice (The Jackson Laboratory Japan, Inc.), OXT-ires-Cre mice (a kind gift from Dr. Brad Lowell, Beth Israel Deaconess Medical Center & Harvard Medical School; (Wu *et al*., 2012)), AVP-Cre mice (a kind gift from Dr. Michihiro Mieda, Kanazawa University; (Mieda *et al*., 2015)) were obtained. Cholecystokinin A receptor floxed mice (CCK-AR^flox/flox^) were generated as described below. All mice were group-housed under controlled temperature (22.5 ± 2°C), humidity (55 ± 10%), and a 12-hour light/dark cycle (lights on 07:30–19:30). Standard laboratory chow (CE-2, CLEA Japan, Tokyo, Japan) and water were available *ad libitum*. Purchased mice were allowed to acclimate to the facility for at least one week before any procedures. All experiments were performed on male mice aged 8–20 weeks.

### Generation of Phox2b-Cre; CCK-AR^flox/flox^ mice

CCK-AR^flox/flox^ mice were generated from frozen sperm obtained from the Helmholtz Zentrum Muenchen - German Research Center for Environmental Health (Ali Khan *et al*., 2023). Mice regenerated from this sperm initially contained a promoter-driven neomycin selection cassette. To remove this cassette, the mice were crossed with C57BL/6-Tg(CAG-flpe)16Ito mice (Kanki *et al*., 2006), which were provided by RIKEN BRC through the National BioResource Project of the MEXT/AMED, Japan. The resulting heterozygous offspring were subsequently intercrossed to generate homozygous CCK-AR^flox/flox^ mice. To generate mice with a selective deletion of the *Cckar* gene in Phox2b-expressing neurons, homozygous CCK-AR^flox/flox^ mice were crossed with Phox2b-Cre mice. The resulting Phox2b-Cre; CCK-AR^flox/flox^ mice were used for experiments. Male littermates (CCK-AR^flox/flox^ mice) lacking the Phox2b-Cre transgene were used as controls.

### Materials

CCK octapeptide (CCK-8; sulfated from 26-33) from Peptide Institute (Osaka, Japan), the CCK-AR antagonist, devazepide from Sigma-Aldrich, the non-selective β-adrenergic receptor antagonist, propranolol hydrochloride from Fujifilm Wako (Osaka, Japan), the β2-adrenergic receptor antagonist, butoxamine hydrochloride from Sigma-Aldrich (St. Louis, MO, USA), the β3-adrenergic receptor antagonist, L-748,337 from Santa Cruz Biotechnology (Dallas, TX, USA), clozapine N-oxide (CNO) from Cayman Chemical (Ann Arbor, MI, USA), the oxytocin receptor antagonist, (d(CH2)_1/5_, Tyr(Me)_2_, Orn^8^)-oxytocin (OVT) from Bachem (Torrance, CA, USA), the vasopressin V1a receptor antagonist SR 49059 from MedChemExpress (Monmouth Junction, NJ, USA) and the vasopressin V1b receptor antagonist SSR-149415 from MedChemExpress (Monmouth Junction, NJ, USA) were obtained.

Two adeno-associated virus (AAV) vectors were purchased from Addgene (Watertown, MA, USA): AAV2-hSyn-DIO-hM4Di-mCherry (Cat# 44362; titer 2.2 × 10^13^ vg/mL) and AAV9-hSyn-FLEX-GCaMP6s (Cat# 50459; 2.5 × 10^13^ vg/mL). An AAV vector designed to express GCaMP6s under the control of the oxytocin promoter (AAV9-OTp-GCaMP6s; titer 2.91 × 10^13^ vg/mL) was a kind gift from Dr. Kazunari Miyamichi (RIKEN Center for Biosystems Dynamics Research; (Yaguchi *et al*., 2023).

### Subdiaphragmatic vagotomy

Bilateral subdiaphragmatic vagotomy was performed as previously described (Iwasaki *et al*., 2015). In brief, a midline incision was made to provide wide exposure of the upper abdominal cavity in C57BL/6J male mice thar were anesthetized via I.P. injection of a three-drug combination (MMB anesthetic), consisting of medetomidine (0.75 mg/kg; Nippon Zenyaku Kogyo, Koriyama, Japan), midazolam (4.0 mg/kg; Maruishi Pharmaceutical, Osaka, Japan), and butorphanol (5.0 mg/kg; Meiji Seika Pharma, Tokyo, Japan). The bilateral subdiaphragmatic trunks of vagal nerves along the esophagus were exposed and cut. In the sham operation group, these vagal trunks were exposed but not cut. To facilitate recovery following surgery, mice were administered atipamezole (0.75 mg/kg, I.P.; Nippon Zenyaku Kogyo) and kept on a heating pad set to 38°C until ambulatory. Vagotomized and sham-operated mice were maintained on a nutritionally complete liquid diet Ensure-H (Abbott Japan, Tokyo, Japan). One to two weeks after the operation, the experiments of thermoregulation were performed. Successful subdiaphragmatic vagotomy was confirmed by an increase in stomach weight.

### Interscapular BAT sympathectomy (iBAT-Sx)

Bilateral sympathectomy of the interscapular brown adipose tissue (iBAT-Sx) was performed. A small incision was made, and the iBAT pad was gently separated from the underlying muscle in C57BL/6J male mice anesthetized with the previously described MMB anesthetic. The five nerve bundles innervating each lobe of the iBAT were carefully isolated and transected. For sham-operated control mice, the nerve bundles were identified and gently manipulated but kept intact. The incision was then sutured, and atipamezole at 0.75 mg/kg was administered to promote recovery. The mice were allowed to recover for a minimum of 10 days before subsequent experiments.

### Stereotactic surgery for DREADD inhibition, fiber photometry, and I.C.V. injection

Mice were anesthetized via I.P. injection of the MMB anesthetic, then head-fixed in a stereotaxic frame (Narishige, Tokyo, Japan). According to a mouse brain atlas of Franklin and Paxinos (Franklin & Paxinos, 2007), AAV was microinjected and either an optical fiber or a stainless steel cannula was implanted into the target region. After surgery, atipamezole was administered to promote recovery.

AAV microinjections were performed as follows. For chemogenetic experiments, OXT-ires-Cre or AVP-Cre male mice were used. AAV2-hSyn-DIO-hM4Di-mCherry (1.0×10^12^ vg/mL) was injected bilaterally into the paraventricular nucleus of the hypothalamus (PVH; 800 nL per site). For fiber photometry experiments, C57BL/6J male mice were used. AAV9-OTp-GCaMP6s (2.91×10^12^ vg/mL; 400 nL) was injected unilaterally into the PVH. The coordinates for the PVH were (in mm from bregma): AP –0.75, ML ± 0.25, DV –4.75.

For implantation of optical fibers or stainless steel cannulas, an optical fiber (400 µm core, 0.50 NA; RWD Life Science, Shenzhen, China) was implanted directly above the PVH injection site immediately following AAV9-OTp-GCaMP6s delivery (DV –4.65) for fiber photometry. All implants were secured to the skull using dental cement. For intracerebroventricular (I.C.V.) drug administration, C57BL/6J male mice were used. A stainless-steel guide cannula (26-gauge; RWD Life Science, Shenzhen, China) was implanted unilaterally into the right lateral ventricle at the following coordinates (in mm from bregma): AP –0.25, ML 1.00, DV –2.50.

### Measurements of rectal temperature

On the day of the experiment, mice were fasted from 09:00 (ZT 1.5) to prevent food intake-induced thermogenesis. At 14:00 (ZT 6.5), a time point when body temperature is at its circadian minimum, a baseline rectal temperature was measured using a flexible rectal probe connected to a BAT-12 Multipurpose Thermometer (Physitemp Instruments, Clifton, NJ, USA). Immediately after the baseline measurement, I.P. saline or CCK-8 at 8 µg/kg was administered, and rectal temperature was measured at 1, 2, and 3 hours post-injection.

In peripheral antagonist experiments, antagonists were administered I.P. 30 injection before CCK-8: devazepide at 0.2 mg/kg (dissolved in 2% DMSO and 10% Tween 80 in saline), propranolol at 10 mg/kg in saline, butoxamine at 5 mg/kg in saline, L-748,337 at 5 mg/kg (dissolved in 2% DMSO in saline), or their respective vehicles. For chemogenetic inhibition, CNO at 1 mg/kg (dissolved in 1% DMSO in saline) or its vehicle was administered I.P. 60 minutes before the injection of CCK-8 or saline. In I.C.V. antagonist experiments, antagonists were administered I.C.V. injection 60 minutes before CCK-8. These included the OXTR antagonist OVT (3.7 nmol in 2 µL of saline) or a mixture of the V1a and V1b antagonists (0.1 nmol SR 49059 and 0.1 nmol SSR-149415 dissolved in 2 µL of 0.2% DMSO in saline). The corresponding vehicle was also administered I.C.V.

Data were analyzed by calculating the incremental area under the curve (IAUC) for the change in rectal temperature from baseline over the 3-hour period following CCK-8 or saline injection. Values below baseline were treated as negative components in the cumulative sum.

### Electrophysiology

For *in vivo* electrophysiological recordings, mice were anesthetized by I.P. injection with wither a mixture of urethane (0.7 g/kg) and α-chloralose (0.06 g/kg) for afferent nerve recordings, or with urethane alone (1.2 g/kg) for efferent nerve recordings. Rectal temperature was maintained at 35.0 ± 0.5°C using a temperature-controlled heating pad throughout the experiments. Polyethylene catheters were placed in a bronchus for oxygen-enriched air supply, the jugular vein for intravenous (I.V.) drug administration, and the carotid artery for monitoring arterial blood pressure.

Gastric afferent vagal afferent nerve activity and efferent iBAT sympathetic nerve activity (iBAT-SNA) were recorded as previously described (Tanida *et al*., 2018; Kuda *et al*., 2019). In gastric vagal afferent nerve activity recordings, a midline abdominal incision was made to identify the gastric branch of the left vagus nerve along the esophagus. The nerve was then carefully dissected and cut centrally to the recording site. In iBAT-SNA recordings, a suprascapular skin incision was made to expose the sympathetic nerve bundles innervating the iBAT. A nerve filament was dissected and cut peripherally to the recording site. In both preparations, the dissected nerve filament was placed on a pair of bipolar stainless-steel wire electrodes. The nerve and electrodes were then covered with silicone gel (Shinohara Chemicals Co., Kagawa, Japan) for electrical insulation and to prevent dehydration.

The recording electrodes were connected to a differential amplifier. The raw electrical signal was filtered with a band-pass of 100–1000 Hz and monitored on an oscilloscope. The processed nerve activity and the arterial blood pressure signal were simultaneously sampled using a PowerLab data acquisition system (8/30; ADInstruments, Bella Vista, NSW, Australia) and recorded on a computer.

After the recorded signals had stabilized, saline as vehicle or CCK-8 at 8 µg/kg was administered as an I.V. bolus injection at a volume of 1 µL/g body weight. The raw nerve activity was rectified and integrated for analysis. The integrated nerve activity was normalized and expressed as a percentage of the pre-stimulus baseline activity. After the I.V. administration experiment was completed, hexamethonium bromide (20 mg/kg), a sympathetic ganglionic blocker, was administered intravenously after cooling stimulation to check valid measurement of postganglionic nerve signal in the iBAT-SNA. At the end of each experiment, the animal was euthanized, and the post-mortem background noise was recorded. This noise level was then subtracted from the integrated signal to determine the true nerve activity.

### Fiber photometry

Fiber photometry recordings were conducted at least 3 weeks after AAV injection. On the day of the experiments, mice were fasted for at least 3 hours prior to the experiment, and recordings were performed between ZT 3–9. After a 15-minute baseline recording period, all mice received an I.P. injection of the antagonist, followed by an I.P. injection of CCK-8 one minute later. Excitation light (473 nm; Dragon Lasers, Changchun, China) was delivered through the implanted optical fiber to record Ca^2+^ signals from PVH^OXT^ neurons. The resulting GCaMP6s fluorescence was collected and filtered (500–550 nm) using an integrated fluorescence mini cube (Doric Lenses, Quebec, QC, Canada). The laser power at the fiber tip was adjusted to approximately 20 µW. Signals were acquired at 1 kHz, passed through a 20 Hz low-pass filter, and downsampled to 5 Hz for analysis using a PowerLab data acquisition system (4sp; ADInstruments, Bella Vista, NSW, Australia).

The change in cytosolic Ca^2+^ concentration from baseline (ΔF/F) was calculated as ΔF/F(%)=100×(F_t_−F_baseline_)/F_baseline_, where F_t_ is the fluorescence at a given time point t, and F_baseline_ is the average fluorescence during a 15-min baseline period immediately preceding the stimulus. To visualize the activity profile around an injection event, peri-event traces were generated by extracting ΔF/F data from –15 min to +60 min relative to the injection. These data were then averaged into 3-minute bins for plotting. The IAUC as calculated as the cumulative sum of the binned ΔF/F values from 0 to 60 min post-injection, with values below baseline treated as negative components.

### Measurements of mRNA expression

Tissues were collected for two distinct experimental aims. First, to assess the knockdown efficiency of the *Cckar* gene in vagal sensory neurons, tissue samples from the hypothalamus, dorsal vagal complex (DVC), nodose ganglion (NG), dorsal root ganglion (DRG), lung, stomach, gallbladder, pancreas, and kidney were collected from CCK-AR^flox/flox^ mice and Phox2b-Cre; CCK-AR^flox/flox^ mice under ad libitum feeding conditions. Second, to investigate the effect of CCK-8 on thermogenic gene expression, iBAT was dissected 1 hour after an I.P. injection of saline or CCK-8 at 8 µg/kg in control mice including sham-operated, iBAT-Sx mice, or SDVx mice. Mice were fasted from 09:00 (ZT 1.5) on the day of the experiment, and the saline or CCK-8 was administered at 14:00 (ZT 6.5). The effect of the β3-adrenergic receptor antagonist L-748,337 was examined by pretreatment at 13:30.

Total RNA was isolated from collected tissues using RNAiso Plus (Takara Bio Inc., Shiga, Japan) according to the manufacturer’s protocol. Subsequently, cDNA was synthesized from DNase-treated total RNA using the ReverTra Ace qPCR RT Master Mix (Toyobo, Osaka, Japan). Real-time PCR was performed using the THUNDERBIRD NEXT SYBR qPCR Mix (Toyobo) on a CFX Connect Real-Time PCR system (Bio-Rad, Hercules, CA, USA). The cycle threshold (Ct) value for each reaction was determined by the system software. Gene expression levels were calculated by first normalizing the Ct value of each target gene to that of the housekeeping gene *36b4*, yielding the ΔCt value (ΔCt = Ct_target_−Ct_36b4_). For *Cckar* expression and basal expression of thermogenic genes, ΔCt values were directly compared between experimental groups. For analyzing the effect of CCK-8 treatment on *Ucp1* and *Pgc1a* expression, relative expression levels were calculated using the comparative Ct (ΔΔCt) method, where results were expressed as fold-change relative to the corresponding saline-injected control group. The following primer sequences were used: 5′-AAGTGACGCTATGCAGCAGT-3′ and 5′-TCACAACCCCAGGGATAAGGA-3′ (*Cckar*); 5′-CTGTGTGTCAGAGTGGATTGGA -3′ and 5′-CTGTGTGTCAGAGTGGATTGGA-3′ (*Ucp1*); 5′-CGGAAATCATATCCAACCAG-3′ and 5′-TGAGAACCGCTAGCAAGTTTG-3′ (*Pgc1a*); 5′-GACCTGGAAGTCCAACTACT-3′ and 5′-CTGCTGCATCTGCTTGGAGC-3′ (*36b4)*.

## Histochemistry

CCK-8 (8 µg/kg, I.P.) was administered to overnight-fasted mice at ZT 2.5 to examine its effect on c-Fos expression in the hypothalamus. Ninety minutes post-injection, mice were deeply anesthetized and transcardially perfused with Zamboni solution (4% paraformaldehyde and 0.2% picric acid in 0.1 M phosphate buffer, pH 7.4). For histological confirmation of viral vector expression such as hM4Di-mCherry or GCaMP6s, brains were processed using the same perfusion protocol. Following perfusion, brains were post-fixed in the same fixative for 4 hours at 4°C, then cryoprotected by sequential overnight incubation in PBS containing 15% and 30% sucrose, respectively Finally, brains were embedded in O.C.T. compound (Tissue-Tek, #4583; Sakura Finetek, Torrance, CA, USA) and stored at –80°C. Coronal sections with a thickness of 40 µm containing the hypothalamus were cut on a cryostat (Leica Biosystems, Wetzlar, Germany), collected at 120 µm intervals, and mounted onto APS-coated glass slides (Matsunami Glass, Osaka, Japan).

Sections were washed three times in PBS. For c-Fos detection, they were permeabilized with 0.1% Triton X-100 in PBS for 10 minutes. All sections were then blocked for 30 min at room temperature in a solution containing 2% bovine serum albumin and 2% normal goat serum in PBS. Subsequently, sections were incubated overnight at 4°C with the appropriate primary antibodies diluted in the blocking solution. The primary antibodies used were: Rabbit anti-c-Fos (#226 004, 1:1000; Synaptic Systems, Goettingen, Germany), Rabbit anti-Oxytocin (#20068, 1:2000; Immunostar, Hudson, WI, USA), Guinea pig anti-Vasopressin (#T-5048, 1:5000; Peninsula Laboratories, San Carlos, CA, USA), Rabbit anti-DsRed (for mCherry; #632496, 1:500; Clontech, Mountain View, CA, USA), and Chicken anti-mCherry (#MCHERRY, 1:300; Aves Labs, Davis, CA, USA). After primary antibody incubation, sections were washed with PBS and incubated for 30 min at room temperature with the appropriate Alexa Fluor-conjugated secondary antibodies (1:500; Thermo Fisher Scientific, Waltham, MA, USA), including goat anti-rabbit Alexa Fluor 488 (#A-11008), goat anti-guinea pig Alexa Fluor 488 (#A-11073), goat anti-rabbit Alexa Fluor 594 (#A-11012), and goat anti-chicken Alexa Fluor 594 (#A-11042). Finally, sections were washed extensively in PBS and coverslipped using Fluoromount (#S3023; Dako, Carpinteria, CA, USA).

Fluorescent images were captured using a DP75 digital camera controlled by cellSens V4.3 imaging software (Evident, Tokyo, Japan), mounted on a microscope with a 20× objective lens (NA 0.80). Images were subsequently processed for brightness and contrast using Affinity Photo (Serif, Nottingham, UK). For quantitative analysis, the number of c-Fos-positive neurons within the PVH (bregma –0.58 to –1.06 mm) was manually counted from at least three sections per animal.

### Statistical analysis

All data are presented as mean ± S.D. unless otherwise stated. Prior to analysis, all data sets were tested for normality using the Shapiro-Wilk test and for homogeneity of variances using the Brown-Forsythe test. As all data met these assumptions, parametric statistical tests were used. Statistical comparisons between two groups were made using a two-tailed unpaired t-test. Comparisons among three or more groups were performed using one-way or two-way analysis of variance (ANOVA), depending on the experimental design. When a significant main effect or interaction was detected by ANOVA, post hoc multiple comparisons were conducted using Tukey’s, Dunnett’s, or Bonferroni’s test, as specified in the figure legends. All statistical analyses were performed using GraphPad Prism 10 (GraphPad Software, San Diego, CA, USA). A p-value < 0.05 was considered statistically significant.

## Results

### The thermogenic effect of exogenous CCK-8 requires CCK-A receptors expressed in vagal sensory nerves

Peripheral administration of CCK-8 has previously been reported to increase local temperature intrascapular brown adipose tissue (iBAT) (Yamazaki *et al*., 2019; Wang *et al*., 2019). However, its effect on core body temperature has remained unclear. To investigate this, we sequentially measured rectal temperature following I.P. administration of CCK-8 in conscious C57BL/6J mice. I.P. administration of CCK-8 at 1, 4, 8, or 16 μg/kg increased rectal temperature in a dose-dependent manner, peaking at 1 hour and lasting for 2 hours (Fig. 1A and B). To determine the involvement of CCK receptors, we pretreated mice with the CCK-A receptor (CCK-AR) antagonist devazepide (DVZ; 0.2 mg/kg). DVZ pretreatment completely abolished the CCK-8-induced thermogenesis (Fig. 1C–E). Surgical subdiaphragmatic vagotomy also eliminated the thermogenic response to CCK-8 (Fig. 1F–H), suggesting an important role for the vagus nerve.

**Figure 1.**
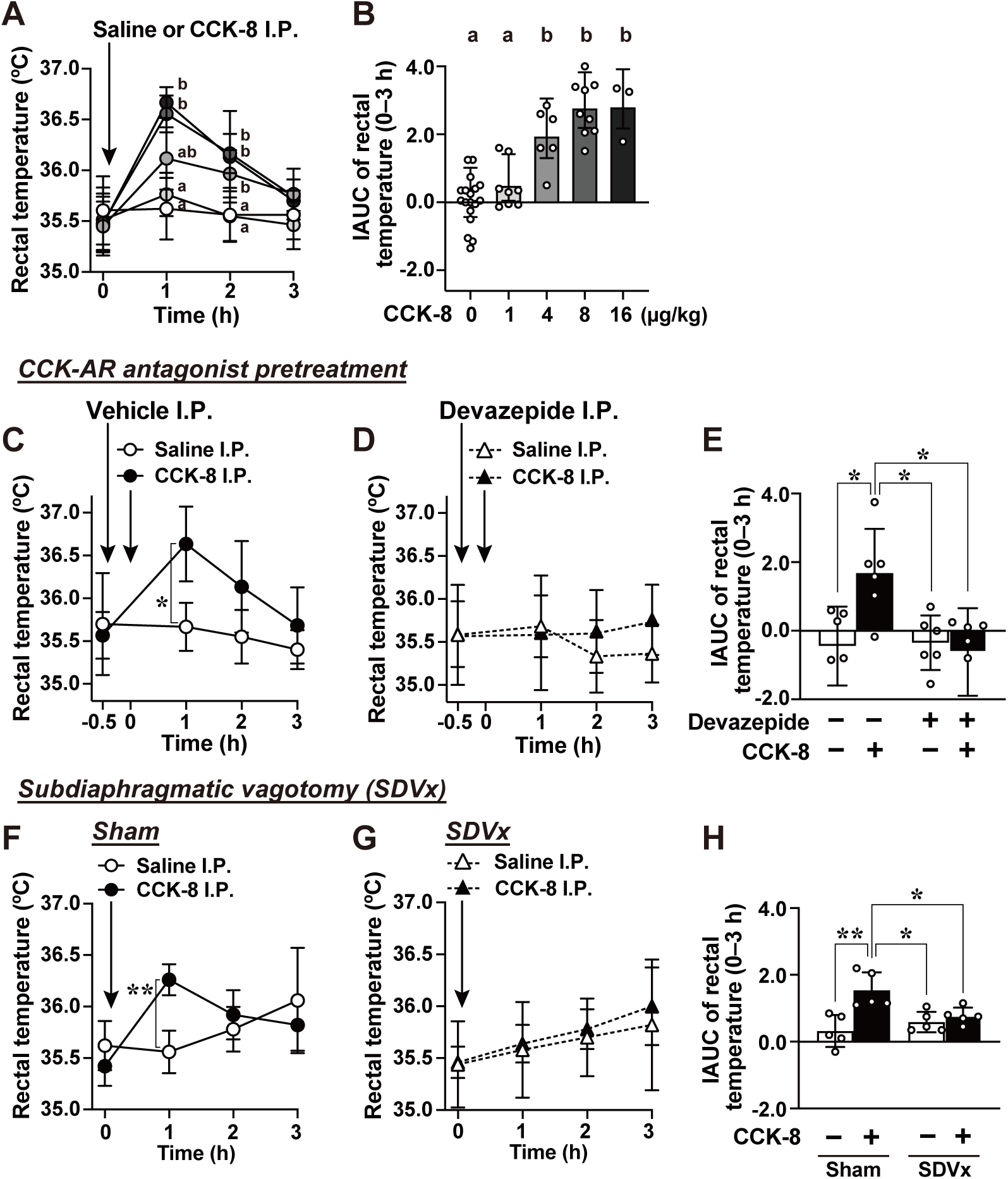
Intraperitoneal injection of CCK-8 increases rectal temperature via CCK-A receptors and vagal sensory nerves. ***A and B***, Time-dependent changes in rectal temperature following intraperitoneal (I.P.) injection of different doses of CCK-8 in male C57BL/6J mice (*A*). Incremental area under the curve (IAUC) for rectal temperature during the 3-hour period following CCK administration (*B*). n = 3–18. Different letters indicate pL<L0.05 by two-way ANOVA followed by Bonferroni’s test in *A*, and different letters indicate pL<L0.05 one-way ANOVA followed by Tukey’s test in B. ***C–E***, Effect of pre-treatment with vehicle or CCK-A receptor antagonist Devazepide on CCK-8-induced rectal temperature elevation. Vehicle (*C*) or Devazepide at 0.2 mg/kg (*D*) was administered intraperitoneally 30 min before I.P. injection of CCK-8 at 8 µg/kg. The results are shown as IAUC (*E*). “Devazepide, –” indicates pre-treatment with vehicle, and “CCK-8, –” indicates injection of saline. ***F–H***, changes in rectal temperature following I.P. injection of CCK-8 at 8 µg/kg in sham-operated mice (*F*) or subdiaphragmatic vagotomized (SDVx) mice (*G*), with the corresponding IAUC (*H*). n = 5–6. In *C*, *D*, *F* and G, **p < 0.01, *p < 0.05 by two-way ANOVA followed by Bonferroni’s test. In *E* and *H*, **p < 0.01, *p < 0.05 by one-way ANOVA followed by Tukey’s test.

To further examine the requirement of CCK-AR expression in vagal sensory neurons, we generated Phox2b-Cre; CCK-AR^flox/flox^ mice (Fig. 2A). Phox2b-Cre mice have been reported to exhibit predominant expression of Cre recombinase in vagal sensory neurons as well as in their central target, the nucleus tractus solitarius (NTS) of the brainstem (Scott *et al*., 2011). Consistent with this report, our Phox2b-Cre; CCK-AR^flox/flox^ mice also showed a ∼98% reduction in *Cckar* mRNA in the nodose ganglion and ∼84% in the NTS, with no significant changes in other tissues (Fig. 2B–D). Next, we performed electrophysiological recordings of vagal afferent nerves in the ventral gastric branch using Phox2b-Cre; CCK-AR^flox/flox^ mice and their littermate controls (CCK-AR^flox/flox^) to assess responses to exogenous CCK-8 administered intravenously. Intravenous injection of CCK-8 at 8 μg/kg significantly increased the firing frequency of vagal afferents in control mice for 0.25–8 minutes and in Phox2b-Cre; CCK-AR^flox/flox^ mice for 0.75–2 minutes (vs. 0 min in each group, p < 0.05, two-way ANOVA with Dunnett’s post hoc test; Fig. 2E). However, the response in Phox2b-Cre; CCK-AR^flox/flox^ mice was markedly attenuated compared with controls (0.25–6 min; Fig. 2E). Subsequently, we assessed the thermogenic response to I.P. CCK-8 administration. Intraperitoneal CCK-8 administration caused a robust rise in rectal temperature in control mice, but this response was completely absent in Phox2b-Cre; CCK-AR^flox/flox^ mice, suggesting a potential involvement of their reduced vagal afferent activity (Fig. 2F and G).

**Figure 2.**
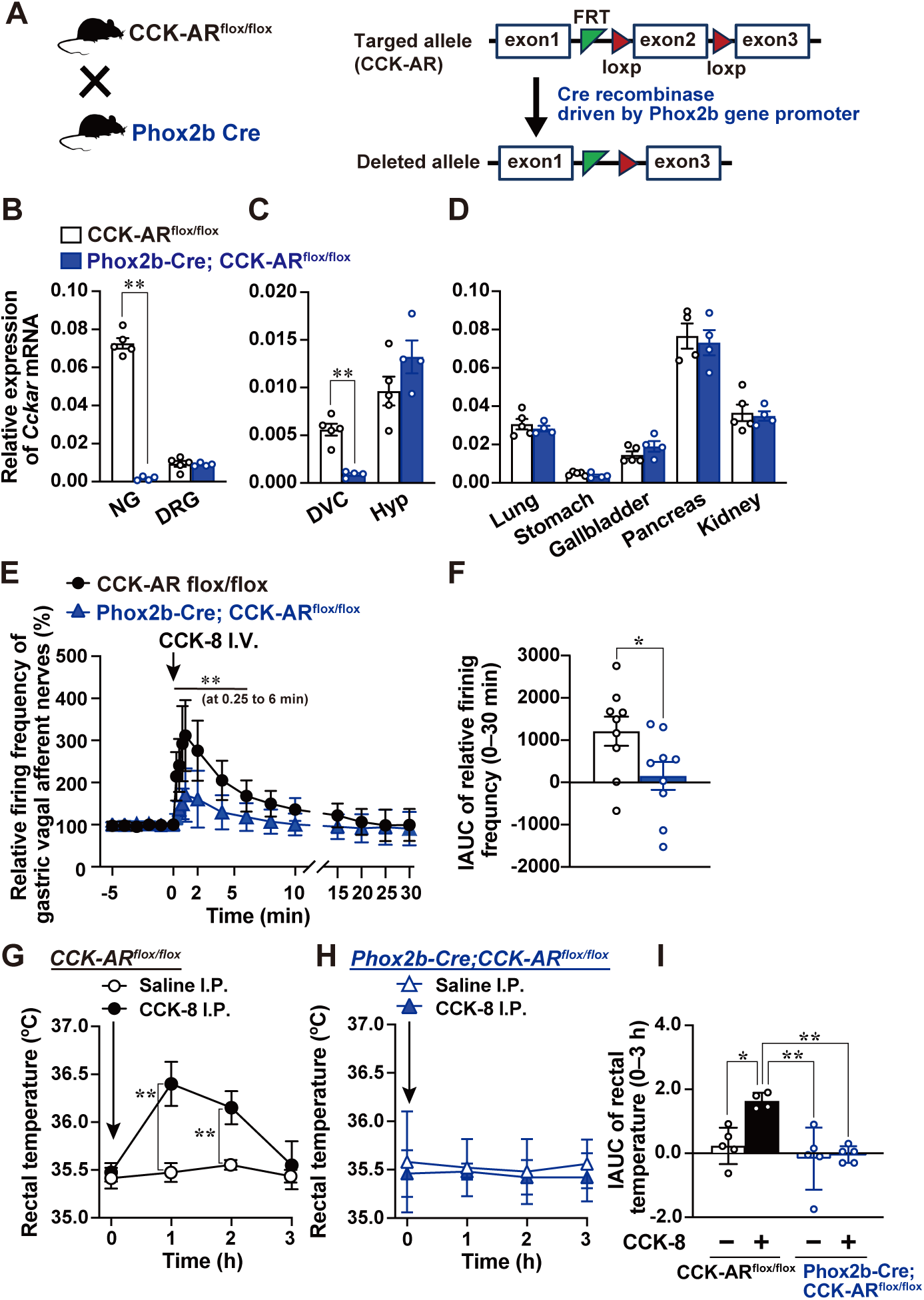
I.P. CCK-8 fails to activate vagal afferents and increase rectal temperature in Phox2b-Cre; CCK-AR^flox/flox^ mice. ***A***, Breeding scheme and schematic illustration of Phox2b-Cre-dependent deletion of CCK-AR. ***B*–*D***, CCK-AR mRNA levels (relative to 36b4) in various tissues from male CCK-AR^flox/flox^ versus Phox2b-Cre; CCK-AR^flox/flox^ mice. Autonomic nerves (nodose ganglion; NG, dorsal root ganglion; DRG), central nerves (dorsal vagal complex; DVC, hypothalamus; Hyp), and peripheral organs were analyzed. n = 4–5. **p < 0.01 by unpaired t-test in *B–D*. ***E and F***, Time-course changes in the relative firing frequency of vagal afferent nerves in the ventral gastric branch following intravenous (I.V.) administration of CCK-8 at 8 µg/kg in CCK-AR^flox/flox^ and Phox2b-Cre; CCK-AR^flox/flox^ mice (*E*), with the corresponding IAUC (*F*). n = 9. **p < 0.01 by two-way ANOVA followed by Bonferroni’s test between groups in E. ***G–I***, Changes in rectal temperature after I.P. injection of CCK-8 at 8 µg/kg in CCK-AR^flox/flox^ (*G*) or Phox2b-Cre; CCK-AR^flox/flox^ mice (*H*), with the corresponding IAUC (*I*). “CCK-8, –” indicates I.P. injection of saline. n = 4–5. **p < 0.01 by two-way ANOVA followed by Bonferroni’s test between groups in G and H. **p < 0.01, *p < 0.05 by one-way ANOVA followed by Tukey’s test in I.

### Thermogenic response to I.P. CCK-8 is mediated by β3-adrenergic receptor signaling

Adrenergic receptor signaling, particularly through β-adrenergic receptors, is well known to mediate thermogenic responses. To assess the involvement of β-adrenergic receptors in CCK-8-induced thermogenesis, we first pretreated mice with the non-selective β-adrenergic receptor antagonist propranolol (10 mg/kg, I.P.). This pretreatment completely abolished the thermogenic response to CCK-8 (Fig. 3A–C). While β1-adrenergic receptors are primarily associated with cardiovascular regulation, β2- and β3-adrenergic receptors are more closely linked to thermogenic functions. In this study, pretreatment with the β3-adrenergic receptor antagonist L-748,337 (5 mg/kg, I.P.) nearly completely suppressed the thermogenic effect of CCK-8 (Fig. 3D–F). In contrast, pretreatment with the β2-adrenergic receptor antagonist butoxamine (5 mg/kg, I.P.) failed to inhibit this response (Fig. 3G–I). These findings indicate that CCK-8-induced thermogenesis is specifically mediated by β3-adrenergic receptor signaling.

**Figure 3.**
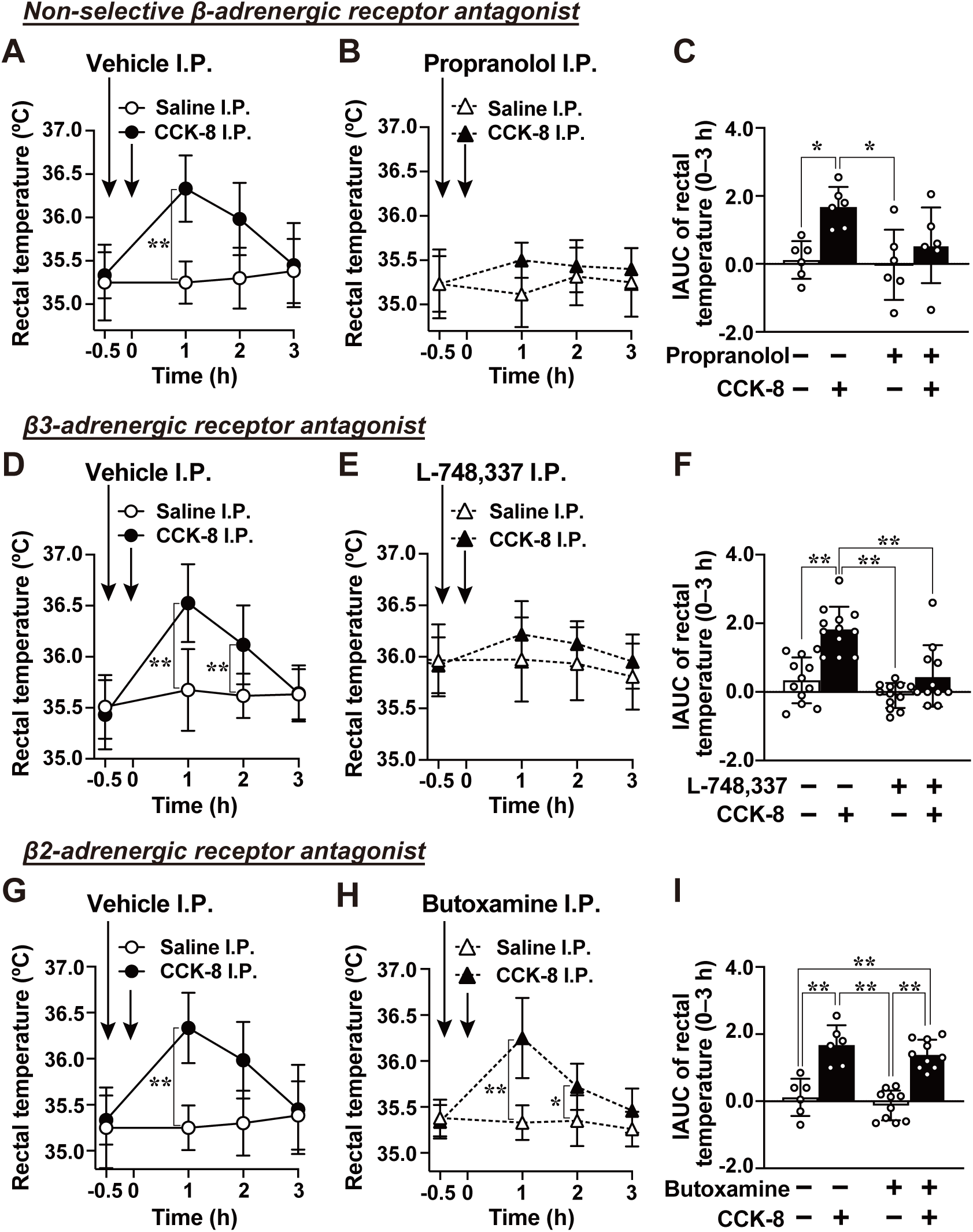
Theβ3-adrenergic, but not the β2-adrenergic, receptor is essential for CCK-induced thermogenesis. ***A*–*I***, Effects of pre-treatment with non-selective, β3-, and β2-adrenergic receptor antagonists on CCK-induced rectal temperature elevation. Vehicle (*A*, *D*, *G*), non-selective β-adrenergic receptor antagonist; propranolol (10 mg/kg, *B*), β3-adrenergic receptor antagonist; L-748,337 (5 mg/kg, *E*), or β2-adrenergic receptor antagonist; butoxamine (5 mg/kg, *H*) was administered intraperitoneally 30 minutes prior to I.P. injection of CCK-8 at 8 µg/kg. The corresponding IAUCs are shown in *C*, *F*, and *I*. In the figure, “–” in each antagonist row indicates vehicle pretreatment, and “CCK-8, –” indicates I.P. injection of saline. n = 6–12. **p < 0.01, *p < 0.05 by two-way ANOVA followed by Bonferroni’s test vs. saline group in *A, B, D, E, G*, and *H*. **p < 0.01, *p < 0.05 by one-way ANOVA followed by Tukey’s test in *C*, *F*, and *I*.

### Exogenous CCK-8 activates iBAT sympathetic nerve activity via vagal afferents

β3-adrenergic receptors are abundantly expressed in iBAT, where noradrenaline released from sympathetic nerve terminals innervating iBAT activates these receptors to induce thermogenesis (Bartness *et al*., 2010). To investigate whether CCK-8 influences this pathway, we performed electrophysiological recordings of sympathetic nerve activity (SNA) innervating the iBAT in C57BL/6J mice. Intravenous administration of CCK-8 at 8 µg/kg induced a biphasic increase in iBAT-SNA: an initial rapid elevation within 0.5–1 minute that returned to baseline, followed by a gradual rise beginning approximately 40 minutes post-injection and sustained for up to 60 minutes (Fig. 4A and B). Both the early and late phases were significantly greater in CCK-8–treated mice compared with saline-treated mice, as confirmed by incremental area under the curve (IAUC) analysis (Fig. 4C and D). Notably, these responses were completely abolished in subdiaphragmatic vagotomized mice, indicating that CCK-8 activates iBAT-SNA in a biphasic manner via vagal sensory nerves.

**Figure 4.**
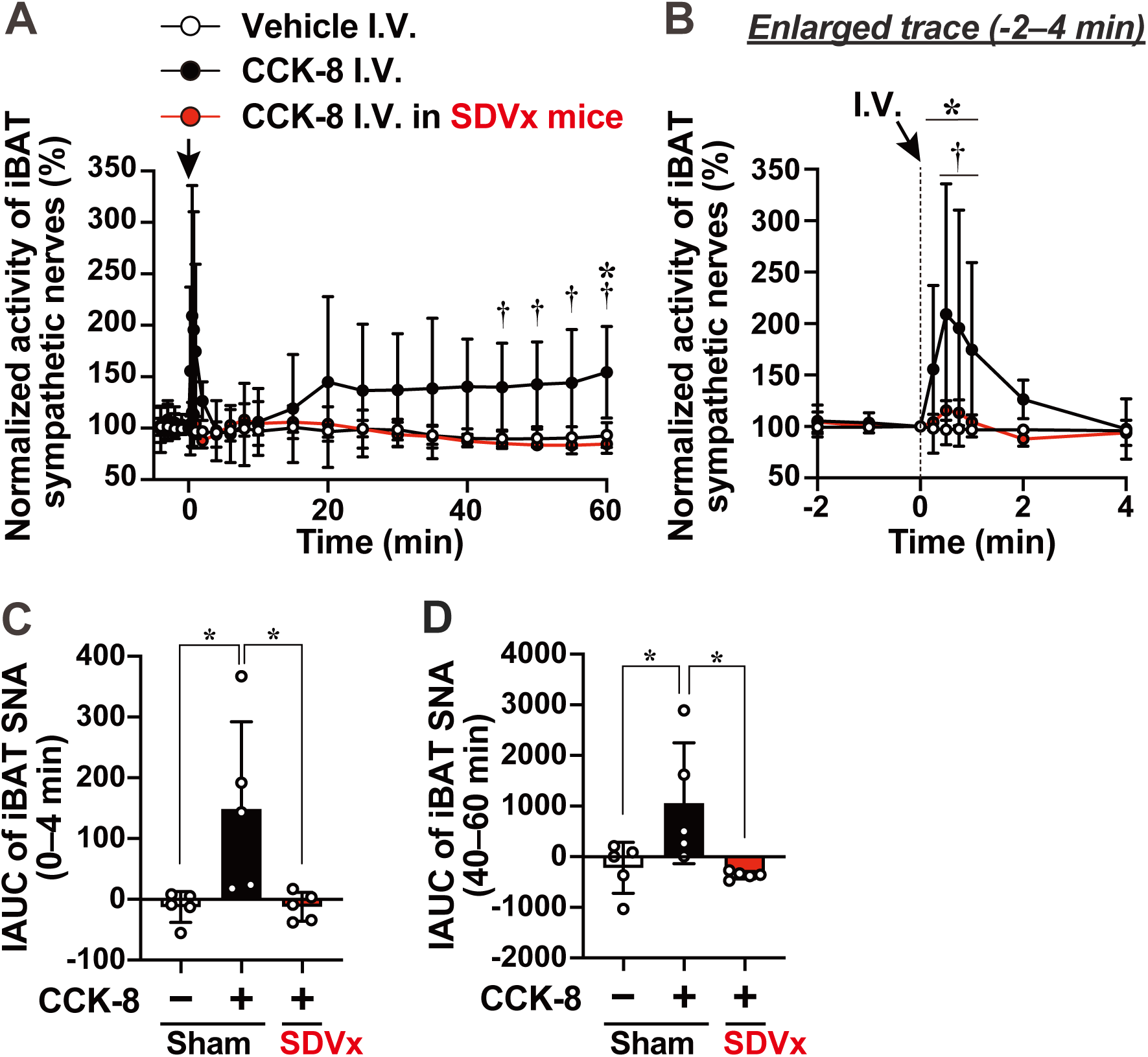
I.P. CCK-8 activates sympathetic nerves innervating intrascapular brown adipose tissue via vagal sensory nerves. ***A***, Time-course of intrascapular brown adipose tissue sympathetic nerve activity (iBAT-SNA) recorded from 4 minutes before to 60 minutes after intravenous (I.V.) injection of CCK-8 at 8 µg/kg in intact and subdiaphragmatic vagotomized (SDVx) mice. In intact mice, CCK-8 administration induced a biphasic sympathetic response: a rapid increase during the initial 0–4 minutes, followed by a gradual and sustained elevation up to 60 minutes. Both the initial and delayed components of this response were abolished by vagotomy. ***B***, Enlarged trace from –2 to +4 minutes from (*A*) to illustrate the initial rapid response. n = 5. *p < 0.05 (CCK-8 I.V. in intact mice vs. vehicle I.V. in intact mice) and †p < 0.05 (CCK-8 I.V. in intact mice vs. CCK-8 I.V. in SDVx mice) by two-way ANOVA followed by Bonferroni’s test. ***C and D***: IAUC of iBAT-SNA during the early (0–4 min, *C*) and late (40–60 min, *D*) post-injection phases. In the figure, “CCK-8, –” indicates I.V. injection of saline. *p < 0.05 by one-way ANOVA followed by Tukey’s test.

### iBAT sympathetic nerves are essential for CCK-induced thermogenesis

To determine the role of iBAT-SNA in CCK-induced thermogenesis, we performed surgical sympathectomy of the interscapular BAT (iBAT-Sx). In sham-operated mice, I.P. injection of CCK-8 at 8 µg/kg increased rectal temperature at 1–2 hours post-injection, whereas this response was markedly attenuated in iBAT-Sx mice (Fig. 5A–C). In iBAT-Sx mice harvested 2 weeks after surgery, mRNA levels of *Ucp1* and *Pgc1a*, encoding key thermogenic proteins, were significantly reduced compared with those in sham-operated mice (Fig. 5D and E). This finding not only confirmed the efficacy of the surgical denervation but also suggested that impaired sympathetic input to iBAT contributed to the attenuation of CCK-induced thermogenesis. Sympathetic stimulation of iBAT is known to drive both heat production and the upregulation of thermogenic gene expression (Klingenberg & Huang, 1999; Barbera *et al*., 2001). We therefore examined whether I.P. injection of CCK-8 alters *Ucp1* and *Pgc1*α mRNA expression in iBAT. To evaluate the roles of vagal sensory input, sympathetic outflow, and β3-adrenergic receptor signaling in this response, we performed surgical interventions and pharmacological blockade targeting these pathways. CCK-8 significantly upregulated *Ucp1* and *Pgc1*α mRNA expression at 1 hour post-injection (Fig. 5F–K). This effect was abolished by SDVx (Fig. 5F and I), iBAT-Sx (Fig. 5G and J), and pretreatment with the β3-adrenergic receptor antagonist L-748,337 (Fig. 5H–K). These results indicate that CCK-8 promotes thermogenesis through vagal sensory input and iBAT-sympathetic outflow, involving transcriptional induction of *Ucp1* and *Pgc1a*.

**Figure 5.**
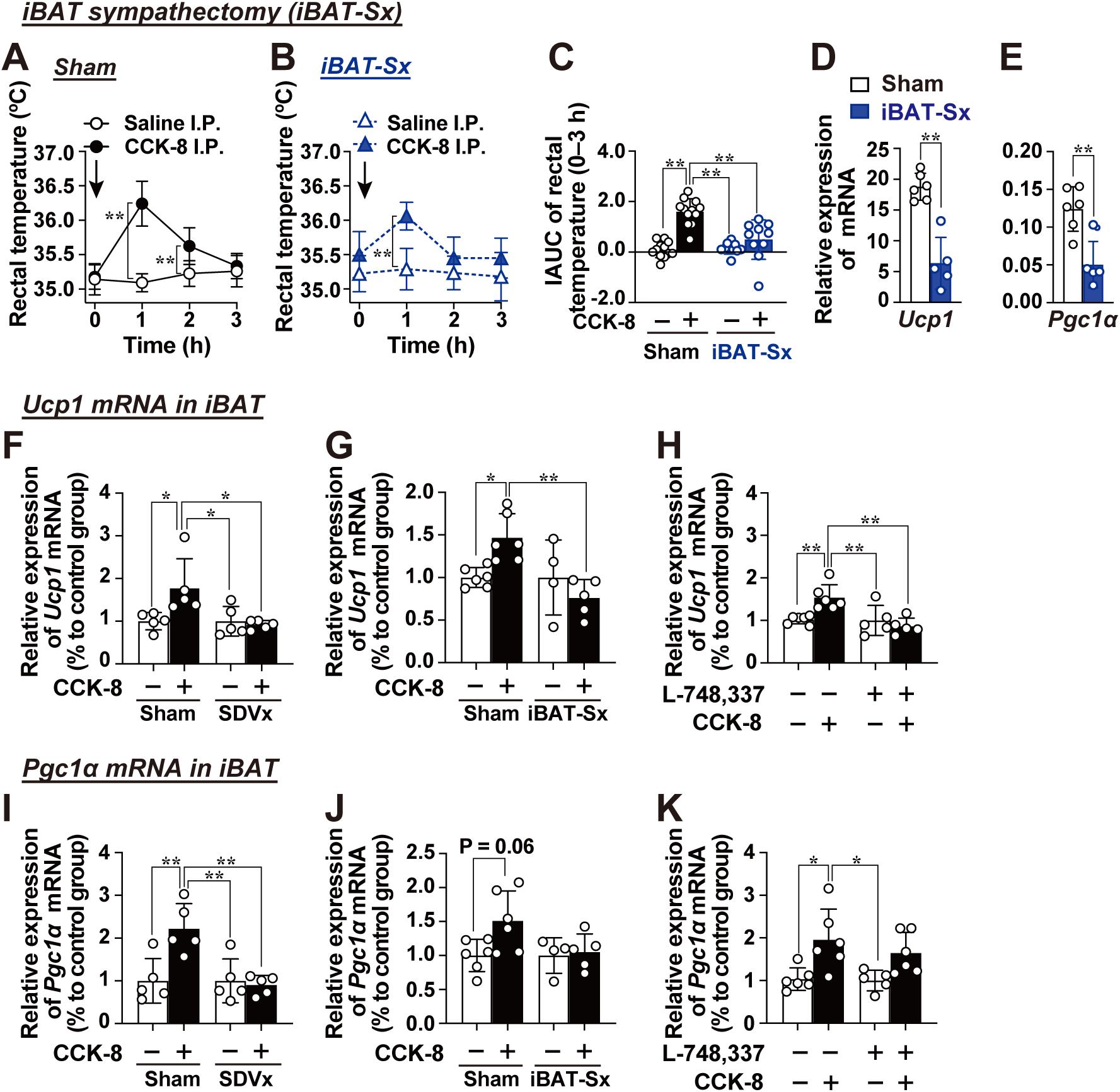
I.P. CCK-8 elevates rectal temperature via non-shivering thermogenesis in iBAT. ***A–C***, Effect of interscapular BAT sympathectomy on CCK-8-induced rectal temperature elevation. Changes in rectal temperature after I.P. CCK-8 at 8 µg/kg in sham-operated (*A*) and iBAT sympathectomized (iBAT-Sx) mice (*B*), with the corresponding IAUC (*C*). n = 10. **p < 0.01 by two-way ANOVA followed by Bonferroni’s test vs. saline group in *A* and *B*. **p < 0.01, *p < 0.05 by one-way ANOVA followed by Tukey’s test in *C*. ***D* and *E***, Basal expression levels of thermogenic genes in iBAT of sham or iBAT-Sx mice. The Y-axis shows ΔCt values of *Ucp1* (*D*) and *Pgc1a* (*E*) mRNA expression, normalized to 36b4. n = 6. **p < 0.01 by unpaired t-test. ***F*–*K***, Relative mRNA expression (ΔΔCt) of *Ucp1* and *Pgc1a* mRNA expression in iBAT 1 hour after CCK-8 I.P. injection. The effects of subdiaphragmatic vagotomy (SDVx; *F* and *I*), iBAT-Sx (*G and J*), and pretreatment with the β3 adrenergic receptor antagonist L-748,337 (*H* and *K*) are shown. n = 4–5. **p < 0.01, *p < 0.05 by one-way ANOVA followed by Tukey’s test. In the figure, “–” in antagonist row indicates vehicle pretreatment, and “CCK-8, –” indicates I.P. injection of saline.

### Peripheral CCK injection rapidly activates PVH oxytocin neurons

To identify the central neuron involved in CCK-induced BAT thermogenesis, we examined the activation of oxytocin neurons in the paraventricular nucleus of the hypothalamus (PVH^OXT^ neurons), which are known to regulate BAT sympathetic output (Freeman & Wellman, 1987; Madden & Morrison, 2009). I.P. administration of CCK-8 at 8 µg/kg significantly increased c-Fos expression in PVH^OXT^ neurons compared to the saline-treated group (Fig. 6A, B). To evaluate the neural dynamics of this activation, we performed *in vivo* fiber photometry recordings in C57BL/6J mice that had received a unilateral PVH-targeted injection of AAV9 expressing GCaMP6s driven by the oxytocin promoter (Fig. 6C and D). Following CCK-8 injection, fluorescence intensity dependent on cytosolic Ca²L levels in PVH^OXT^ neurons exhibited a transient increase, peaking at approximately 3 minutes and gradually returned to baseline within 30 minutes (Fig. E). This Ca²L response was completely suppressed by pretreatment with the CCK-AR antagonist DVZ at 0.2 mg/kg (Fig. 6E and F). These findings demonstrate that peripheral CCK-8 rapidly activates PVH^OXT^ neurons through CCK-AR-dependent mechanisms.

**Figure 6.**
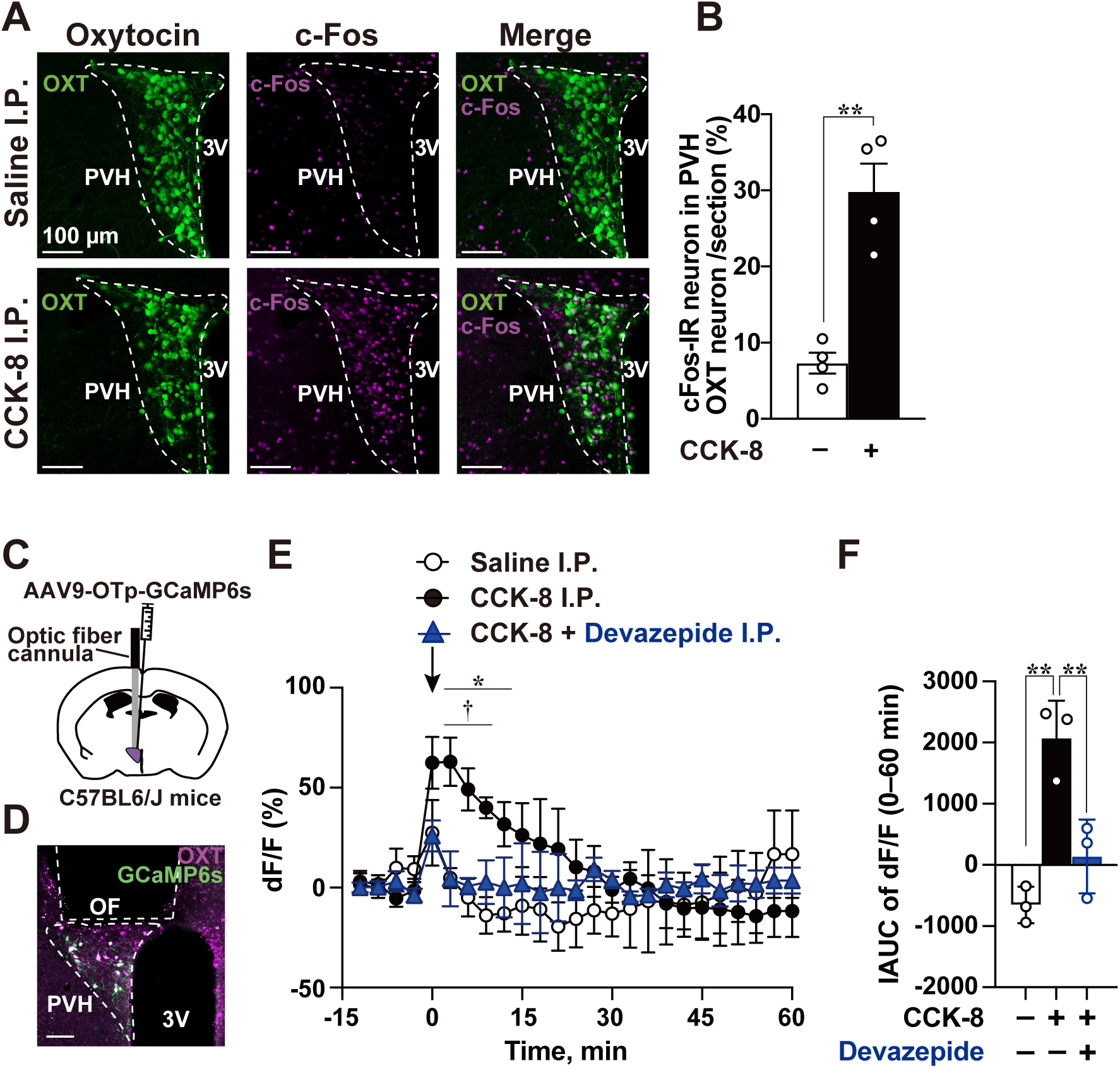
I.P. CCK-8 rapidly activates oxytocin neurons in the paraventricular nucleus of the hypothalamus. ***A***, Representative pseudocoloured images showing oxytocin-immunoreactive neurons (green) and c-Fos-immunoreactive neurons (magenta) in the paraventricular nucleus of the hypothalamus (PVH). Scale bars, 100 µm. 3V, third ventricle. ***B***, Quantification of the percentage of c-Fos-positive PVH^OXT^ neurons in saline- or CCK-8-treated mice. **p < 0.01 by unpaired t-test. n = 4. ***C***, Schematics of the experimental design of fiber photometry. ***D***, Representative coronal section of the PVH showing the location of the optical fiber and expression of the GCaMP6s (green) and oxytocin neuron (magenta). Scale bars, 100 µm. 3V, third ventricle; OF, optic fiber. ***E***, Time-course plots showing the change of cytosolic Ca^2+^ concentration from baseline (ΔF/F) responses averaged over 3-minute intervals (–15 to +60 min) after I.P. injection of saline, CCK-8 at 8 µg/kg, or CCK-8 with CCK-AR antagonist: Devazepide (DVZ) at 0.2Lmg/kg. ***F***, IAUC of the ΔF/F from 15 minutes before to 60 minutes after injection. n = 3. *p < 0.05 (CCK-8 I.P. vs Saline I.P.) and †p < 0.05 (CCK-8 I.P. vs CCK-8 + Devazepide I.P.) by two-way ANOVA followed by Bonferroni’s test in *E*. **p < 0.01 by one-way ANOVA followed by Tukey’s test in *F*. In the figure, “CCK-8, –” indicates I.P. injection of saline and “Devazepide, –” indicates vehicle pretreatment.

### Exogenous CCK drives thermogenesis via PVH^OXT^-OXTR signaling in the brain

Given the potential role of PVH^OXT^ neurons in CCK-8-induced thermogenesis, we used OXT-ires-Cre mice bilaterally injected in the PVH with AAV2-hSyn-DIO-hM4Di-mCherry to selectively inhibit these neurons using an inhibitory DREADD and assess their contribution (Fig. 7A and B). Inhibiting PVH^OXT^ neurons with CNO pretreatment markedly blunted the CCK-8-induced increase in rectal temperature compared with vehicle pretreatment (Fig. 7C–E). To examine whether CCK-induced thermogenesis is specifically regulated by PVH^OXT^ neurons, we focused on another neuronal population within the PVH, the arginine vasopressin (AVP) neurons. AVP acts on overlapping oxytocin and vasopressin receptor systems in the brain (Koshimizu *et al*., 2012). Using a similar inhibitory DREADD system, inhibition of PVH^AVP^ neurons did not reduce CCK-8-induced thermogenesis (Fig. 7G–I). Next, we tested whether this pathway is mediated through oxytocin receptor (OXTR) signaling in the brain. Intracerebroventricular (I.C.V.) injection of the OXTR antagonist OVT markedly blunted the significant thermogenic response to CCK-8 compared with I.C.V. vehicle administration (Fig. 7J–M). Furthermore, to investigate the involvement of central vasopressin V1a and V1b receptors, we co-administered V1a and V1b receptor antagonists I.C.V. prior to CCK-8 I.P. injection. This treatment did not affect the thermogenic response to CCK-8 (Fig. 7N–P). These findings indicate that CCK-8-induced thermogenesis is mediated by PVH^OXT^-OXTR signaling.

**Figure 7.**
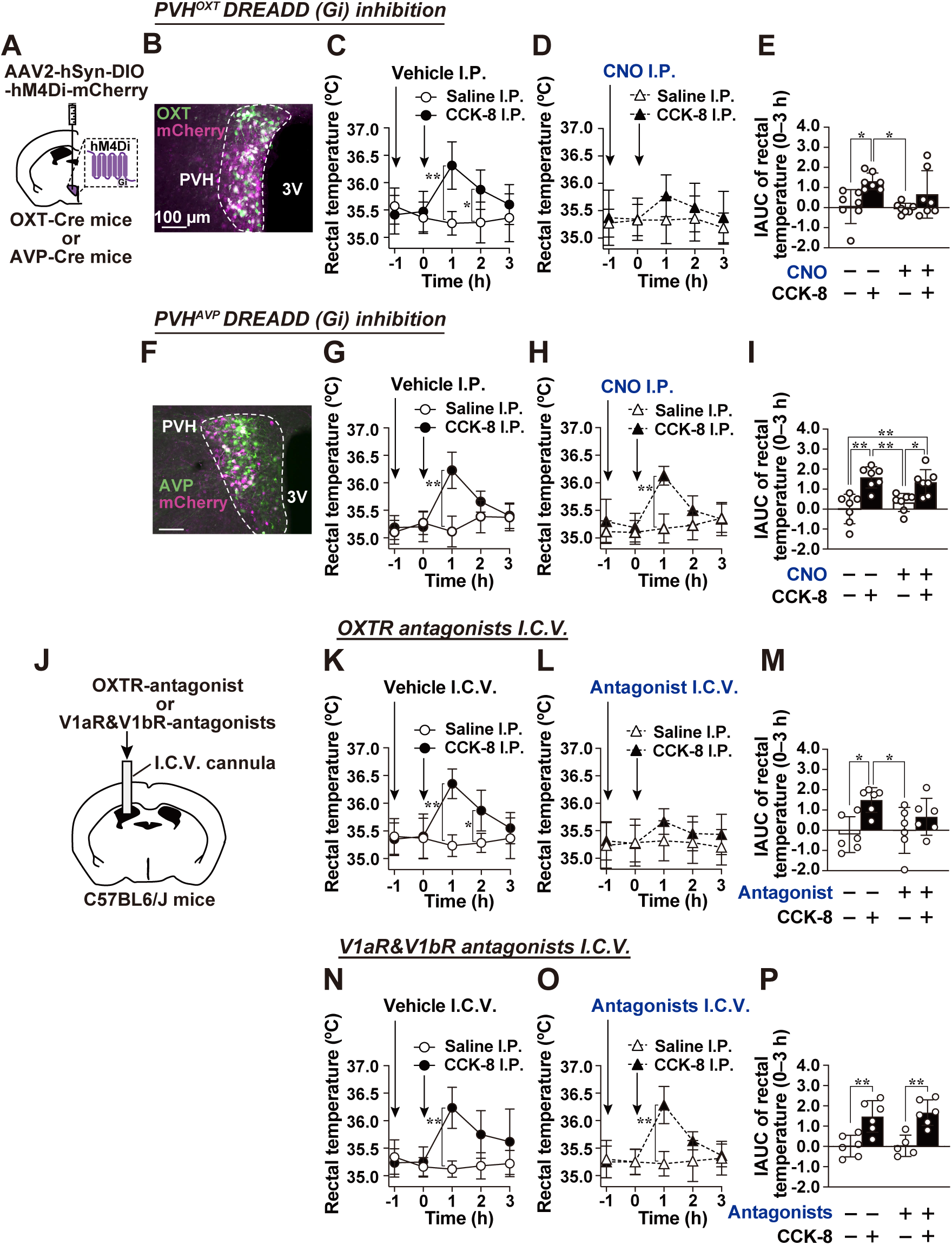
PVH^OXT^ neurons and central OXTR signaling are critical for CCK-induced thermogenesis. ***A***, Experimental design for chemogenetic inhibition using hM4Di-mCherry AAV injection into the PVH of OXT-Cre or AVP-Cre mice. ***B***, A representative coronal section of the PVH showing expression oxytocin (green) and hM4Di-mCherry (magenta). Scale bar, 100 µm. ***C*–*E***, Effect of PVH^OXT^ neuron inhibition on CCK-8-induced rectal temperature increase. OXT-Cre mice expressing hM4Di in PVH^OXT^ neurons received vehicle (I.P.; *C*) or CNO (1 mg/kg, I.P.; *D*) 1 hour prior to I.P. injection of saline or CCK-8 at 8 µg/kg. IAUC from 0 to 3 h is shown in (*E*). n = 7. **p < 0.01, *p < 0.05 by two-way ANOVA followed by Bonferroni’s test vs. saline group in *C* and *D*; one-way ANOVA followed by Tukey’s test in *E*. ***F***, Representative PVH section showing expression AVP (green) and hM4Di-mCherry (magenta). Scale bar, 100 µm. ***G*–*I***, Effect of PVH^AVP^ neuron inhibition on CCK-8-induced rectal temperature increase. AVP-Cre mice expressing hM4Di in PVH^AVP^ neurons received vehicle (*G*) or CNO (*H*) followed by saline or CCK-8, using the same protocol as in *C-E*. IAUC is shown in *I*. n = 6 *p < 0.01, p < 0.05 by two-way ANOVA followed by Bonferroni’s test vs. saline group in *G* and *H*; one-way ANOVA followed by Tukey’s test in *I*. ***J***, Schematics of the I.C.V. cannulation. ***K*–*M***, Effect of I.C.V administration of an OXT receptor antagonist OVT on CCK-8-induced thermogenesis. Vehicle (2 µl, *K*), OVT (3.7 nmol µg in 2 µl, *L*), was administered I.C.V. 1 hour before CCK-8 injection at 8 µg/kg. IAUC is shown in *M*. n = 6. **p < 0.01, *p < 0.05 by two-way ANOVA followed by Bonferroni’s test vs. saline group in *K* and *L*; one-way ANOVA followed by Tukey’s test in *M*. ***N–P***, Effect of I.C.V. co-administration of V1a and V1b receptor antagonists. Vehicle (2 µl, *N*), or the antagonist mixture (0.1 nmol of SR 49059 and 0.1 nmol of SSR-149415 in 2 µl, *O*) was administered I.C.V. 1 hour before CCK-8 injection. IAUC is shown in *P*. n = 6. **p < 0.01, *p < 0.05 by two-way ANOVA followed by Bonferroni’s test vs. saline group in *N* and *O*; one-way ANOVA followed by Tukey’s test in *P*. In the figure, “–” in CNO or each antagonist row indicates vehicle pretreatment, and “CCK-8, –” indicates I.P. injection of saline.

## Discussion

In this study, we aimed to elucidate the physiological role of vagal sensory neurons, which relay information from peripheral organs to the brain, by focusing on the gut hormone CCK. We investigated the thermoregulatory mechanism mediated by CCK’s action on vagal sensory neurons and the associated autonomic reflex pathways. I.P. administration of CCK-8 transiently increased rectal temperature. This effect was markedly attenuated by pretreatment with the CCK-AR antagonist, subdiaphragmatic vagotomy, and CCK-AR knockdown in Phox2b-expressing neurons, primarily targeting vagal sensory neurons. These findings indicate that CCK-AR-expressing vagal sensory neurons are essential for CCK-induced thermogenesis. Furthermore, we found that this afferent input activates sympathetic outflow to the iBAT, leading to the activation of iBAT-projecting sympathetic nerves and rectal temperature elevation via β3-adrenergic receptors. Using fiber photometry in freely moving mice, we also demonstrated that I.P. injection of CCK-8 rapidly and transiently increased cytosolic Ca²L levels in PVH^OXT^ neurons in a CCK-A receptor-dependent manner. Notably, chemogenetic inhibition of PVH^OXT^ neuronal activity or I.C.V. injection of an OXTR antagonist substantially blunted CCK-induced thermogenesis. Collectively, these results reveal a novel neural circuit underlying CCK-induced thermogenesis, comprising CCK-AR-expressing vagal sensory neurons (afferent input), PVH^OXT^ neurons and OXTR signaling (central integrative hub), and iBAT and its innervating sympathetic nerves (efferent output).

Previous studies investigating the thermoregulatory functions of vagal sensory nerves have reported conflicting results, and a unified view has not yet been established. For example, cervical vagus nerve stimulation (VNS) in humans has been shown to activate BAT and increase body temperature (Vijgen *et al*., 2013), whereas VNS in rats has been reported to suppress BAT activity and lower body temperature (Madden *et al*., 2017; Larsen *et al*., 2017). In models of inflammation-induced fever, lipopolysaccharide triggers vagus-dependent fever (Simons *et al*., 1998), while interleukin-1β has been shown to induce vagal sensory nerve-dependent hypothermia (Silverman *et al*., 2023). These findings, together with recent single-cell RNA sequencing analyses of vagal sensory neurons that revealed distinct gene expression profiles (Kupari *et al*., 2019; Bai *et al*., 2019), suggest the existence of functionally heterogeneous vagal sensory neuron subpopulations regulating both thermogenesis and heat dissipation. Therefore, to dissect the thermoregulatory role of vagal sensory nerves, it is critical to target specific neuronal subtypes defined by their receptor expression, rather than broadly activating vagal fibers with electrical stimulation. Our study demonstrates that selective activation of CCK-AR-expressing vagal sensory neurons was key to uncovering the afferent–central–efferent pathways mediating CCK-induced thermogenesis. Specifically, we show that CCK activates vagal afferents, which in turn stimulate PVH^OXT^ neurons and engage central OXTR signaling to drive sympathetic outflow to iBAT. Previous studies have independently shown that CCK administration induces thermogenesis in iBAT (Yamazaki *et al*., 2019; Wang *et al*., 2019) and that PVH^OXT^ neurons enhance sympathetic nerve activity to iBAT, promoting thermogenesis (Fukushima *et al*., 2022). Our findings integrate these two lines of evidence and establish a functional connection between CCK-AR-expressing vagal sensory neurons, PVH^OXT^ neurons/OXTR signaling, and iBAT sympathetic nerves, forming a coherent neural circuit for CCK-indued thermoregulation.

Endogenously secreted CCK regulates a variety of postprandial responses, including its anorexigenic effect, inhibition of gastric emptying, stimulation of pancreatic exocrine secretion, gallbladder contraction, and suppression of hepatic glucose production, some of which are mediated via vagal sensory nerves (Raybould & Taché, 1988; Cheung *et al*., 2009; Dockray, 2012; Iwasaki & Yada, 2012). Among postprandial physiological processes, diet-induced thermogenesis (DIT) is of particular interest, raising the possibility that the CCK-induced thermogenesis and its underlying neural mechanism identified in this study may contribute to DIT. Supporting this notion, Blouet and colleagues reported that duodenal lipid infusion in rats increases iBAT temperature, an effect abolished by I.P. administration of a CCK-AR antagonist and by local microinjection of a glutamate N-methyl-D-aspartate (NMDA) receptor antagonist into the NTS, a central target of vagal sensory neurons (Blouet & Schwartz, 2012). Thus, the CCK-induced thermogenic pathway elucidated here may also be driven by endogenously released CCK during the postprandial state. Future studies are warranted to determine whether CCKAR-expressing vagal sensory neurons are activated by meal-derived CCK to promote thermogenesis.

With the global rise in obesity and metabolic disorders such as diabetes, there is growing interest in developing novel intervention strategies to enhance energy expenditure. Among these, DIT, which increases resting metabolic rate after meals, has emerged as an attractive physiological mechanism for enhancing metabolism without pharmacological intervention. However, the neural regulatory mechanisms underlying DIT remain poorly understood. In this study, we elucidated the role of CCK-AR-expressing vagal sensory neurons, PVH^OXT^ neurons, and iBAT-projecting sympathetic nerves in mediating CCK-induced thermogenesis and its underlying mechanism. Interestingly, previous studies have reported that in diet-induced obesity models, CCK-induced activation is attenuated at multiple steps of this pathway, including vagal sensory neurons (Wang *et al*., 2019), PVH^OXT^ neurons (Gruber *et al*., 2023), and BAT thermogenesis (Wang *et al*., 2019), suggesting the development of CCK resistance. Moreover, PVH^OXT^ neurons are critically involved not only in enhancing energy metabolism but also in preventing hyperphagia (Zhang *et al*., 2011; Iwasaki *et al*., 2019; Inada *et al*., 2022). Thus, dysfunction of the CCK-mediated thermogenic pathway identified in this study may contribute to the pathogenesis of obesity and diabetes. Conversely, identifying strategies to preserve or restore this pathway may offer novel approaches for the prevention and treatment of metabolic diseases.

## Additional information

### Data availability statement

The data that support the findings of this study are available from the corresponding author upon reasonable request.

### Author contributions

Y.M., M.K., M.T. and Y.I. developed concept and designed the study. Y.M., M.K., R.K., K.O, M.T. and Y.I. performed experiments and analyzed data. T.W., H.Y. and M.T. developed the novel mouse strain. Y.M., M.K., M.T., and Y.I. prepared figures, interpreted the results of the experiments, and drafted the manuscript. All of the authors edited the final draft. All authors have read and agreed to the published version of the manuscript.

## Acknowledgements

The authors thank Drs. Hirokazu Hirai and Ayumu Konno (Gunma University) for developing the AAV9-OTp-GCaMP6s vector, and Dr. Kazunari Miyamichi (RIKEN Center for Biosystems Dynamics Research) for technical advice on fiber photometry and generously providing the AAV.

